# Plasmacytoid dendritic cells produce type I interferon and reduce viral replication in airway epithelial cells after SARS-CoV-2 infection

**DOI:** 10.1101/2021.05.12.443948

**Authors:** Luisa Cervantes-Barragan, Abigail Vanderheiden, Charlotte J. Royer, Meredith E. Davis-Gardner, Philipp Ralfs, Tatiana Chirkova, Larry J. Anderson, Arash Grakoui, Mehul S. Suthar

**Author notes:** Correspondence: Mehul S. Suthar and Luisa Cervantes-Barragan. Lead contact: Mehul S. Suthar.

## Abstract

Infection with SARS-CoV-2 has caused a pandemic of unprecedented dimensions. SARS-CoV-2 infects airway and lung cells causing viral pneumonia. The importance of type I interferon (IFN) production for the control of SARS-CoV-2 infection is highlighted by the increased severity of COVID-19 in patients with inborn errors of type I IFN response or auto-antibodies against IFN-α. Plasmacytoid dendritic cells (pDCs) are a unique immune cell population specialized in recognizing and controlling viral infections through the production of high concentrations of type I IFN. In this study, we isolated pDCs from healthy donors and showed that pDCs are able to recognize SARS-CoV-2 and rapidly produce large amounts of type I IFN. Sensing of SARS-CoV-2 by pDCs was independent of viral replication since pDCs were also able to recognize UV-inactivated SARS-CoV-2 and produce type I IFN. Transcriptional profiling of SARS-CoV-2 and UV-SARS-CoV-2 stimulated pDCs also showed a rapid type I and III IFN response as well as induction of several chemokines, and the induction of apoptosis in pDCs. Moreover, we modeled SARS-CoV-2 infection in the lung using primary human airway epithelial cells (pHAEs) and showed that co-culture of pDCs with SARS-CoV-2 infected pHAEs induces an antiviral response and upregulation of antigen presentation in pHAE cells. Importantly, the presence of pDCs in the co-culture results in control of SARS-CoV-2 replication in pHAEs. Our study identifies pDCs as one of the key cells that can recognize SARS-CoV-2 infection, produce type I and III IFN and control viral replication in infected cells.

**Importance:** Type I interferons (IFNs) are a major part of the innate immune defense against viral infections. The importance of type I interferon (IFN) production for the control of SARS-CoV-2 infection is highlighted by the increased severity of COVID-19 in patients with defects in the type I IFN response. Interestingly, many cells are not able to produce type I IFN after being infected with SARS-CoV-2 and cannot control viral infection. In this study we show that plasmacytoid dendritic cells are able to recognize SARS-CoV-2 and produce type I IFN, and that pDCs are able to help control viral infection in SARS-CoV-2 infected airway epithelial cells.

## Introduction

Infection with SARS-CoV-2 in humans has caused a pandemic of unprecedented dimensions. In the United States, more than 31 million people have tested positive for infection and more than 550,000 have died as of April 2021. SARS-CoV-2 infects airway epithelial cells and Type 2 pneumocytes causing fever, dry cough, and shortness of breath. While most patients experience mild to moderate disease, some progress to more severe disease, including pneumonia and acute respiratory failure. Severe cases of COVID-19 can result in lung damage, low blood oxygen levels, and death^1,2^.

Type I and III IFNs are a major part of the innate immune defense against viral infections^3^. One characteristic of coronavirus infections including SARS-CoV-2 is the low induction of type I IFN in the host^4–6^. The reduced type I IFN production from infected cells is caused by both lack of recognition of the viral RNA by pathogen recognition receptors, and the IFN antagonist function of several viral non-structural proteins (nsp; as reviewed in^7,8^). Early reports detected low levels of type I IFN in the blood of COVID-19 patients^4^. However, transcriptional analysis of bronchoalveolar lavage (BAL) or peripheral blood of COVID-19 patients detected an early type I IFN signature^9–11^. The importance of type I IFN production for the control of SARS-CoV-2 infection is highlighted by the increased severity of COVID-19 in patients with inborn errors of type I IFN response, as well as patients presenting with auto-antibodies against IFN-α^12,13^. Moreover, IFN-2α treatment of COVID-19 patients with inborn errors of type I IFN response, as well as early administration of IFN-α2 to COVID-19 patients reduced in hospital mortality^14,15^.

Plasmacytoid dendritic cells (pDCs) are a unique immune cell population specialized in recognizing and controlling viral infections through the production of high concentrations of type I IFN^16,17^. pDCs have unique features that enable their unparalleled antiviral response. They express pattern recognition receptors like TLR7 or TLR9^18^, which allow pDCs to sense viral RNA or DNA even in the absence of viral infection or replication^19^. They constitutively express IRF-7, the master transcription factor for interferon (IFN)-α, which enables rapid production and secretion of IFN-α^16,17^. Both human and murine pDCs can sense SARS-CoV and the murine coronavirus MHV-A59 respectively. pDCs can sense these coronaviruses through TLR7 and produce high concentrations of IFN-α^20,21^. pDCs numbers are reduced in the blood of COVID-19 patients^9^. However, increased numbers of pDCs are found in the BAL of mild COVID-19 patients in contrast to severe COVID-19 patients which have reduced numbers of pDCs in BAL^10^. Network analyses of single-cells RNA seq of COVID-19 patients showed an association between apoptosis in pDCs and disease severity^22^. Despite these findings, we still lack studies that address the contribution of pDCs to SARS-CoV-2 control in the airways.

Primary human airway epithelial cell cultures are permissive to SARS-CoV-2 infection, and have been used for both short and long term modelling of infection. Ciliated cells and goblet cells are permissive to infection while SARS-CoV-2 has not been detected on basal and club cells ^23^. Our previous studies have shown that human airway epithelial cell cultures (pHAE) are unable to produce a type I or III IFN response after infection with SARS-CoV-2^24^. Similarly, Lieberman et al observed no induction of interferon stimulated genes (ISGs) after 3 days of SARS-CoV-2 infection in pHAEs ^25^. A delayed type I IFN response is observed in pHAEs after 7 days of infection and positively correlates with the levels of replication of SARS-CoV-2^26^, which is recognized by pHAE cells through MDA-5^27^. Importantly, in contrast to SARS-CoV, SARS-CoV-2 is susceptible to type I and III IFN inhibition^28,29^. Pre- or post-treatment of pHAEs with IFN-β or IFN-λ was able to reduce SARS-CoV-2 replication, suggesting that if an IFN response can be elicited, it is effective in controlling SARS-CoV-2 infection^24^.

In this study, we tested the ability of pDCs to sense SARS-CoV-2. pDCs isolated from healthy donors recognized SARS-CoV-2 and rapidly produced large concentrations of IFN-α. Sensing of SARS-CoV-2 by pDCs was independent of viral replication. We observed that pDCs can recognize UV-inactivated SARS-CoV-2 and produce type I IFN with similar kinetics. Transcriptional profiling of SARS-CoV-2 and UV-SARS-CoV-2 showed a swift type I and III IFN signature after 12h stimulation. Moreover, we observed the induction of several chemokines, pro-inflammatory cytokines and the induction of apoptosis in pDCs. To test if type I IFN production by pDCs was able to control viral replication we modeled SARS-CoV-2 infection in the lung using primary human airway epithelial cells (pHAEs) and showed that co-culture of pDCs with SARS-CoV-2 infected pHAEs induces an antiviral response and upregulation of antigen presentation in pHAE cells. Importantly, the presence of pDCs in the co-culture results in control of SARS-CoV-2 replication in pHAEs. Overall, our study identifies pDCs as a key cells that recognize SARS-CoV-2 infection and produce type I and III IFN that can control viral replication in airway cells.

## Materials and Methods

### Viruses and cells

The infectious clone SARS-CoV-2 (icSARS-CoV-2) was kindly provided to us by Dr. Vineet Menachery (UTMB)^30^. Viral titers were determined by plaque assay or focus-forming assay on VeroE6 cells (ATCC). VeroE6 cells were cultured in complete DMEM medium consisting of 1x DMEM (Corning Cellgro), 10% FBS, 25 mM HEPES Buffer (Corning Cellgro), 2 mM L-glutamine, 1mM sodium pyruvate, 1x Non-essential Amino Acids, and 1x antibiotics. Viral stocks were titered on VeroE6 cells and stored at −80°C until use.

### Plasmacytoid dendritic cell isolation

Deidentified human blood from healthy individuals was collected with the approval of the Internal Review Board (IRB) of Emory University number IRB00045821. PBMCs were obtained by Ficoll-Hypaque density gradient centrifugation. pDCs were further purified by magnetic sorting with human Diamond Plasmacytoid dendritic-cell isolation kit II (Miltenyi Biotec). Purity of cell preparation was assessed by flow cytometry and for all donors was more than 90%.

### Plasmacytoid dendritic cell stimulation

4×10^4^ pDCs were infected with SARS-CoV-2 (MOI=1) or stimulated with medium, UV inactivated SARS-CoV-2 (MOI=1) or CpG A (6ug/ml). After one-hour medium was replaced and cells were incubated at 37°C. After 12hr cell were collected and frozen for RNA isolation and after 24h supernatants were collected and frozen for ELISA measurements.

### Generation of primary human airway epithelial cells

pHAE cultures were generously provided by Dr. C. U. Cotton (Case Western Reserve University) and cultured as described previously^31^. Briefly, bronchial lung specimens were seeded on Transwell Permeable Support Inserts (Costar-Corning) and cultured until confluent. Specimens were then transferred to an air/liquid interface and differentiated for 2-3 weeks. Cultures were maintained in DMEM/Ham’s F-12 medium supplemented with 2% Ultroser G (Pall Corp., France), until all wells had TEER measurements greater than 1000 Ω and deemed ready for use.

### Co-culture of HAE and pDC cells

pDCs were isolated as described above, then stimulated for 3 hours in complete RPMI supplemented with; medium, CpG A (ODN2216 at 6μg/ml), UV inactivated SARS-CoV-2 (MOI=1) at 37°C. During the 3 hour stimulation, differentiated pHAE cultures were infected; the apical side of the pHAE culture was washed 3 times with PBS, then SARS-CoV-2 (MOI=1) was allowed to adsorb for 1 hour at 37°C. After adsorption, the apical side was washed 3 times with PBS to remove excess virus. The basolateral media was then replaced with fresh media containing the stimulated pDCs, medium only, or IFNβ (100 IU/mL). Cells were co-cultured for up to 72 hours, with no disturbance of the basolateral media. To collect viral supernatant, PBS was added to the apical side and incubated for 30 minutes at 37°C.

### Focus-forming Assays

To measure SARS-CoV-2 viral burden, supernatant from infected cells was serially diluted (10-fold dilutions) in serum free RPMI. Virus dilutions were overlaid on VeroE6 cells and incubated with a methylcellulose overlay (0.85% methylcellulose in 2X DMEM) for 48 hours at 37°C. Methylcellulose was removed, cells were then fixed with 2%-PFA and permeabilized with 0.1% bovine serum albumin (BSA)-Saponin in PBS. Permeabilized cell monolayers were incubated with an anti-SARS-CoV-2 spike protein primary antibody (provided by Jens Wrammert, Emory University) conjugated to biotin for 2 hours at room temperature (RT), then washed 3 times. Cells were incubated with an avidin-HRP conjugated secondary antibody for 1 hour at RT. Foci were visualized using True Blue HRP substrate and imaged on an ELISPOT reader (CTL Analyzers)^24^.

### IFN-α ELISA

Human IFN-α concentration in cell-culture supernatants was measured by enzyme-linked immunosorbent assay (ELISA; PBL Biomedical Laboratories, Piscataway, NJ) according to the manufacturer’s instructions.

### Quantitative PCR analysis

pHAE cultures were lysed by adding RNA lysis buffer directly to the apical layer for >5 minutes, then lysate was collected in Eppendorf tubes. RNA was extracted using the Zymo Quick-RNA MiniPrep kit (VWR, R1055) according to the manufacturers protocol, then reverse transcribed into cDNA using a high-capacity cDNA reverse transcription kit (Thermo Fisher, 43-688-13). RNA levels were quantified using the IDT Prime Time Gene Expression Master Mix, and Taqman gene expression Primer/Probe sets. All qPCR was performed in 384-well plates and run on a QuantStudio5 qPCR system. Viral RNA was quantified using SARS-CoV-2 Rdrp specific primers and probes as described in Vanderheiden et al. The following Taqman Primer/Probe sets were used in this analysis: Gapdh (Hs02758991_g1), IFIT2 (Hs01922738_s1), IFIH1 (Hs00223420_m1). C_T_ values were normalized to the reference gene GAPDH and represented as fold change over mock.

### RNA-Sequencing

pDCs were centrifuged and lysis buffer was added to the pellet. RNA was extracted using the RNAeasy microkit (Qiagen) following manufacturer’s instructions. RNA from pHAE cells was extracted as described above, and mRNA sequencing libraries were prepared by the Yerkes Genomics Core using the Clontech SMART-Seq v4 kit. Barcoding and sequencing primers were added using a NexteraXT library kit, and validated by microelectrophoresis. Libraries were sequenced on an Illumina NovaSeq 6000 and mapped to the human reference genome 38 using DNASTAR software. Viral genes were mapped to the FDAARGOS_983 strain of the 2019-nCoV/USA-WA1/2020 SARS-CoV-2 isolate. Reads were normalized and differentially expressed genes were analyzed using DESeq2 (Bioconductor). Gene set enrichment analysis was performed using the software provide by the Broad Institute and the MSigDB database. The raw data of all RNA sequencing will be deposited into the Gene Expression Omnibus (GEO) repository and the accession number will be available following acceptance of this manuscript.

### Statistical analysis

Statistical analyses were performed using GraphPad Prism 8, ggplot2 R package, and GSEA software. Statistical significance was determined as P value of 0.05 using Student’s t test or a one-way analysis of variance (ANOVA). All comparisons were made between treatment or infection conditions with time point-matched, uninfected and untreated controls.

## Results

### Plasmacytoid Dendritic cells (pDCs) recognize SARS-CoV-2 and produce type I IFN

pDCs are uniquely poised to respond to pathogen infection and produce large amounts of type I IFNs. However, the role of pDCs in responding to SARS-CoV-2 is not well understood. To investigate this, pDCs were isolated from blood of healthy donors and infected with SARS-CoV-2. CpG A (ODN2216 at 6μg/ml) was used as control. Following SARS-CoV-2 infection, pDCs produced over 3500 pg/ml of IFN-α as early as 24 hours post-infection (**Figure 1A**). To determine if virus replication is required for pDCs to respond to SARS-CoV-2, we stimulated pDCs with either live SARS-CoV-2 or UV inactivated SARS-CoV-2 (UV-SARS-CoV-2) at equivalent MOIs and measured IFN-α production. pDCs maintained IFN-α production after stimulation with UV-SARS-CoV-2 as compared to live SARS-CoV-2, demonstrating that virus replication is not required by pDCs for sensing SARS-CoV-2 and to produce type I IFN.

**Figure 1.**
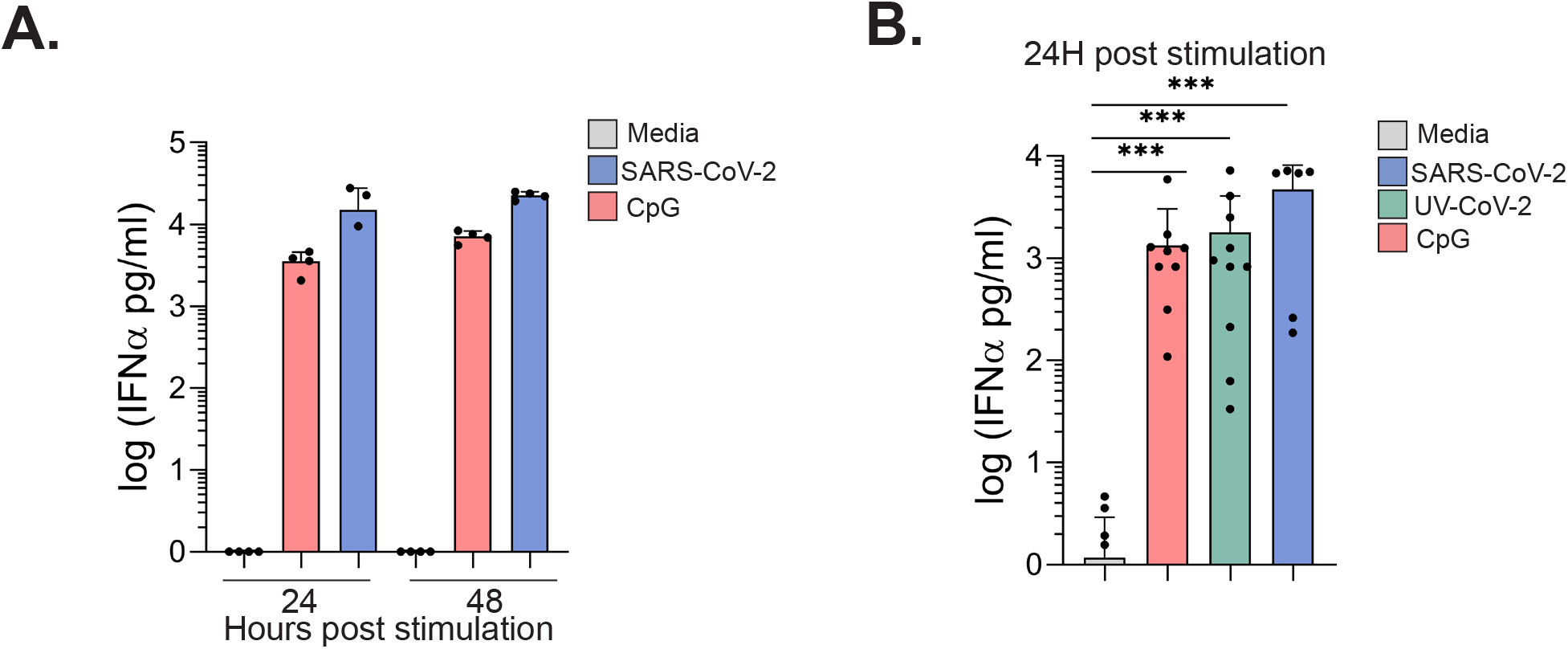
Plasmacytoid dendritic cells (pDCs) recognize SARS-CoV-2 and produce IFN-α. A) human pDCs were mock infected (media), infected with SARS-CoV-2 (MOI=1) or stimulated with CpG A (6μg/ml) for 24 or 48 hr. B) human pDCs were stimulated with medium, SARS-CoV-2, UV inactivated SARS-CoV-2 or CpG A for 24h. At the indicated timepoints supernatants were collected and IFN-α was measured by ELISA. Dots represent pDCs from single donors, bars depict means. Graphs are representative of 3 independent experiments. Data was analyzed using one-way ANOVA ***, P<0.001.

### Transcriptional profile of SARS-CoV-2 stimulated pDCs shows type I and type III IFN and apoptosis signature

Besides their ability to produce type I IFN, pDCs are able to produce several proinflammatory cytokines and chemokines as well as type III IFN ^17^. To determine the response of pDCs to SARS-CoV-2, and identify if there are different expression patterns when pDCs are stimulated with SARS-CoV-2 vs. UV-SARS-CoV-2 or CpG, pDCs were stimulated with SARS-CoV-2, UV-SARS-CoV-2 and CpG, RNA was isolated 12h post stimulation and bulk RNAseq performed. pDCs stimulated with either SARS-CoV-2 or UV-SARS-CoV-2 expressed high levels of IFN-β, IFN-α as well as type III IFN: IFN-λ1 and IFN-λ3 and IFN-ω. SARS-CoV-2 and UV-SARS-CoV-2 stimulated pDCs also highly expressed several chemokines such as CXCL10 (IP-10) and CXCL11 (IP-9) CCL7 (MCP3), CCL8 (MCP2), CCL2 (MCP-1) which recruit monocytes and T cells to sites of inflammation, as well as promote T cell adhesion^32^. Finally, pDCs also upregulated IDO expression which has been associated with pDC-dependent induction of T regulatory cells (Tregs)^33,34^ (**Figure 2A-C**). When comparing stimulation of pDCs with SARS-CoV-2 to pDCs stimulated with either UV-SARS-CoV-2 or CpG A we confirmed that the type I and III IFN gene expression was similar in all three groups (**Figure 2C and D**). However, there were some DEGs were present only each of the conditions, Interestingly, we also observed an increase in the apoptosis signature in SARS-CoV-2 activated pDCs when compared to medium stimulated pDCs suggesting pDCs may die after activation with SARS-CoV-2 and type I and III IFN production (**Figure 2E**). These results confirm that viral replication is not needed for a high production of type I and type III IFN by pDCs after sensing SARS-CoV-2 infection, and that stimulation with SARS-CoV-2 may increase the apoptosis signature in pDCs.

**Figure 2.**
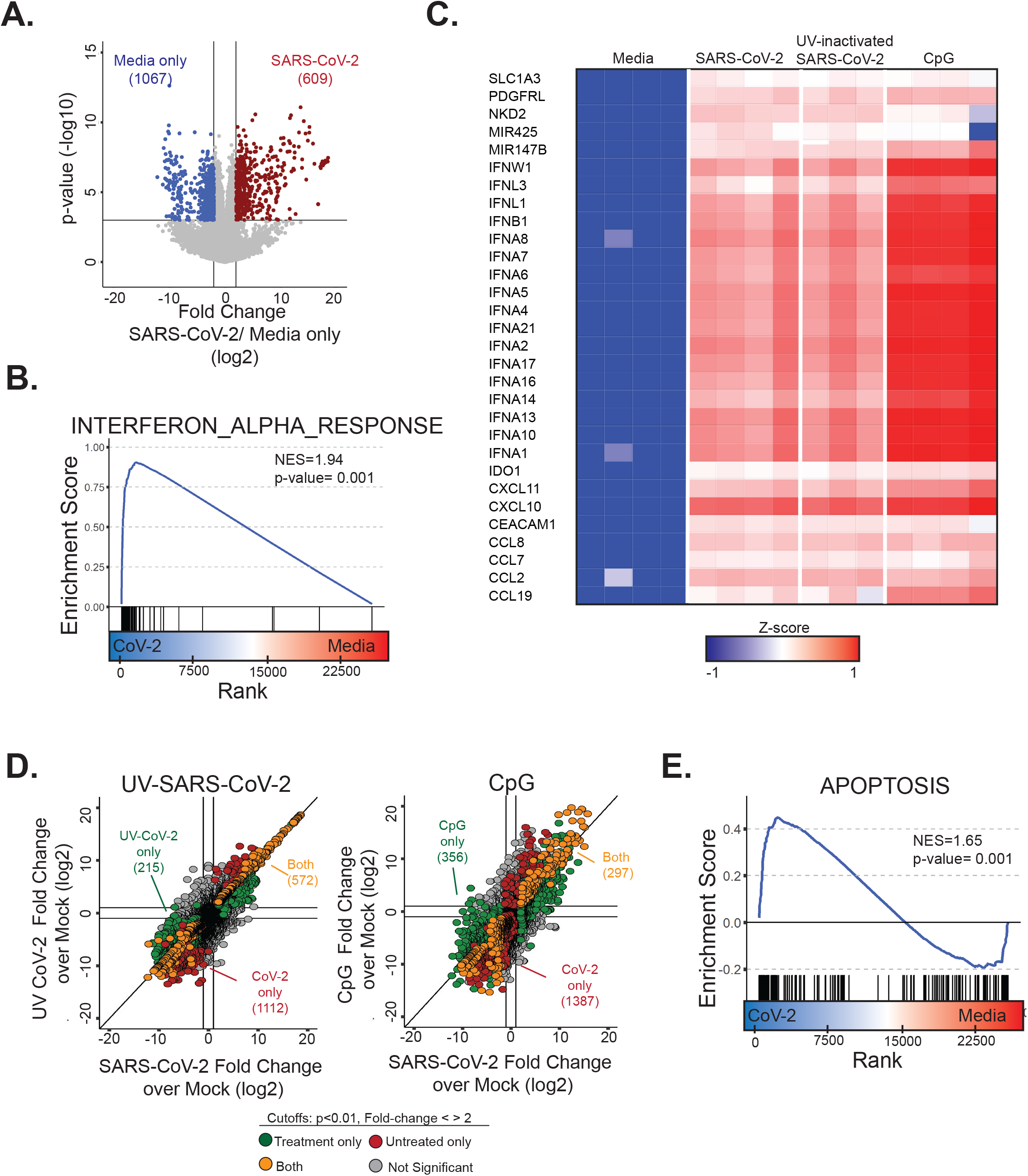
pDC transcriptional profile shows a type I and type II IFN signature after SARS-CoV-2 stimulation. Human pDCs were infected with SARS-CoV-2 (MOI=1) or stimulated with UV-SARS-CoV-2 (MOI=1), CpG A (6μg/ml) or medium. After 12h cells were collected for bulk-RNA-Seq analysis. A) Volcano plot showing all differentially regulated genes (DEG) between media and SARS-CoV-2 infected pDCs in red and blue. B) GSEA plot showing enrichment in interferon-alpha-response genes after SARS-CoV-2 activation in pDCs. C) Heatmap illustrating the z-scores for the 30 most differentially upregulated genes in SARS-CoV-2 infected pDCs. D) Fold-change vs fold-change plot of SARS-CoV-2 vs medium and UV-SARS-CoV-2 vs medium showing in green the DEG in SARS-UV-CoV-2 only, in red the DEG in SARS-CoV-2 only and in orange the DEG in both conditions and fold-change vs fold-change plot of SARS-CoV-2 vs medium and CpG A vs medium showing in green the DEG in CpG only, in red the DEG in SARS-CoV-2 only and in orange the DEG in both conditions. E) GSEA plot showing enrichment in Apoptosis genes in medium vs SARS-CoV-2 stimulated pDCs.

### pDCs protect primary human airway epithelial (pHAE) cells from SARS-CoV-2 infection

Since pDCs are able to produce type I and III IFN after SARS-CoV-2 stimulation, we next investigated if pDC-derived-IFN is able to control SARS-CoV-2 replication and protect lung epithelial cells. To test this, we utilized pHAE cells isolated from the bronchial or tracheal region of healthy donors. These cells are cultured in an air-liquid interface to create a polarized, pseudostratified epithelial layer that models the critical features of the human respiratory tract, such as cilium movement and mucus production^31^. We have previously shown that pHAE cells are susceptible to SARS-CoV-2 infection, and respond by producing a variety of inflammatory cytokines^35^. However, SARS-CoV-2 infection does not induce the production of type I or type III IFN by pHAE cells. To test if pDCs are able to control viral infection in pHAE cells, we developed a co-culture system. pHAE cells were infected with SARS-CoV-2 at MOI=1 for 1 hour. In parallel, pDCs were stimulated with UV-SARS-CoV-2, CpG A, or left untreated. After 3 hours, pDCs were washed and placed in the bottom well of the SARS-CoV-2 infected-pHAE transwell culture (**Figure 3A**). Addition of recombinant IFN-β, UV-SARS-CoV-2 and CpG A to the bottom wells in the absence of pDCs was used as control. UV-SARS-CoV-2 stimulated pDCs and CpG stimulated pDCs were able to strongly reduce replication and virus production of SARS-CoV-2 in pHAE cells. Co-culture with stimulated pDCs reduced infectious SARS-CoV-2 production (as measured by focus-forming assay) from the apical surface of pHAE cells by 1000-fold, as well as viral RNA (100-fold reduction) at 72h post infection (**Figure 3B and C**). Importantly, the protective effect was mediated by pDCs, since addition of CpG or UV-SARS-CoV-2 alone to the basolateral side didn’t reduce viral replication. Interestingly, unstimulated pDCs were also able to reduce the viral load, suggesting that infected pHAE cells are able to activate a type I IFN response in pDCs that is rapid enough to decrease viral replication within 72 hours (**Figure 3B and C**). As seen in our previous study, addition of IFN-β to the bottom well reduced viral replication, confirming its protective effect against SARS-CoV-2 and suggesting pDCs act through type I IFN production. Indeed, when we analyzed the transcriptional profile of infected pHAEs co-cultured with pDCs we observed the upregulation of several interferon-stimulated genes (ISGs) such as IFIH1 and IFIT2 (**Figure 3D**).

**Figure 3.**
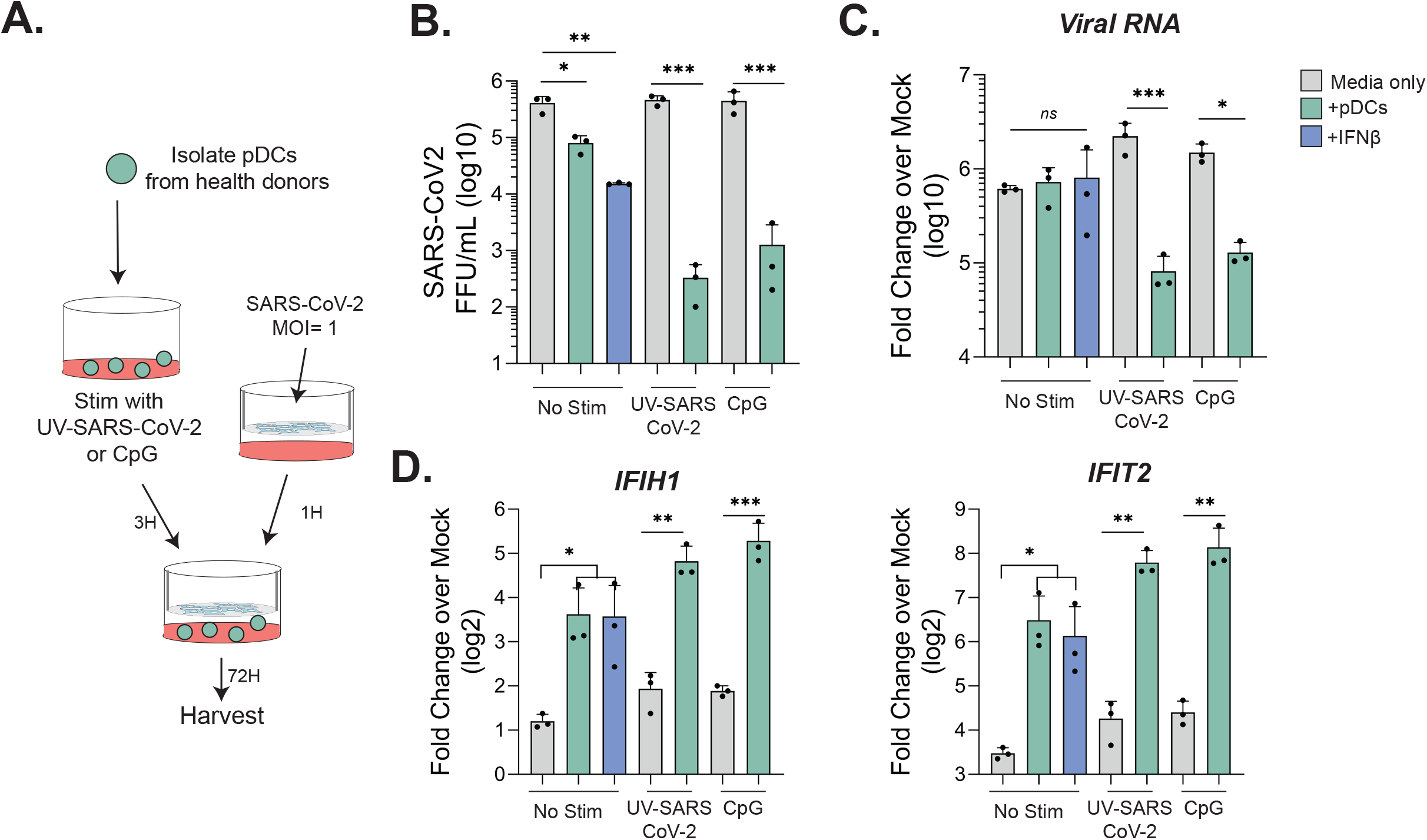
pDCs reduce SARS-CoV-2 replication in pHAE cultures. A) Experimental schematic. B) pDCs were stimulated for 3 hours in complete RPMI supplemented with; medium, CpG A (ODN2216 at 6μg/ml), or UV inactivated SARS-CoV-2 (MOI=1) at 37°C. In parallel, differentiated pHAE cultures were infected with SARS-CoV-2 (MOI=1). After 3 hours the basolateral media was replaced with fresh media containing the stimulated pDCs, medium only, or IFNβ (100 IU/mL). After 72 hours, the apical side was washed and supernatant was collected and pHAE cells were harvested and RNA extracted. B) Viral titers in the apical side, C) Total viral RNA in the cells and D) IFIH1 and IFIT2 expression in pHAEs in the different conditions tested. Dots represent single replicates and bars represent the means. Data was analyzed using one-way ANOVA. *, P<0.05; **, P<0.01; ***, P<0.001.

### Activated pDCs induce an antiviral profile in SARS-CoV-2 pHAE infected cells

We next evaluated how the presence of activated pDCs changes the transcriptional profile of SARS-CoV-2 infected pHAE cells. To this end, we performed mRNAseq analysis of pHAE cells infected with SARS-CoV-2 and co-cultured with pDCs, UV-SARS-CoV-2 activated pDCs, and IFN-β or media as controls. First, analysis at 72 hrs post infection showed a reduction in viral read counts spanning the whole viral genome when infected pHAEs cells were co-cultured with activated pDCs. Interestingly, co-culture with activated pDCs was more efficient than addition of IFN-β in reducing viral RNAs, which is likely due to the concerted action of type I IFN (IFN-β and IFN-α) and type III IFN (IFN-λ) produced by pDCs (Figure 4A).

**Figure 4.**
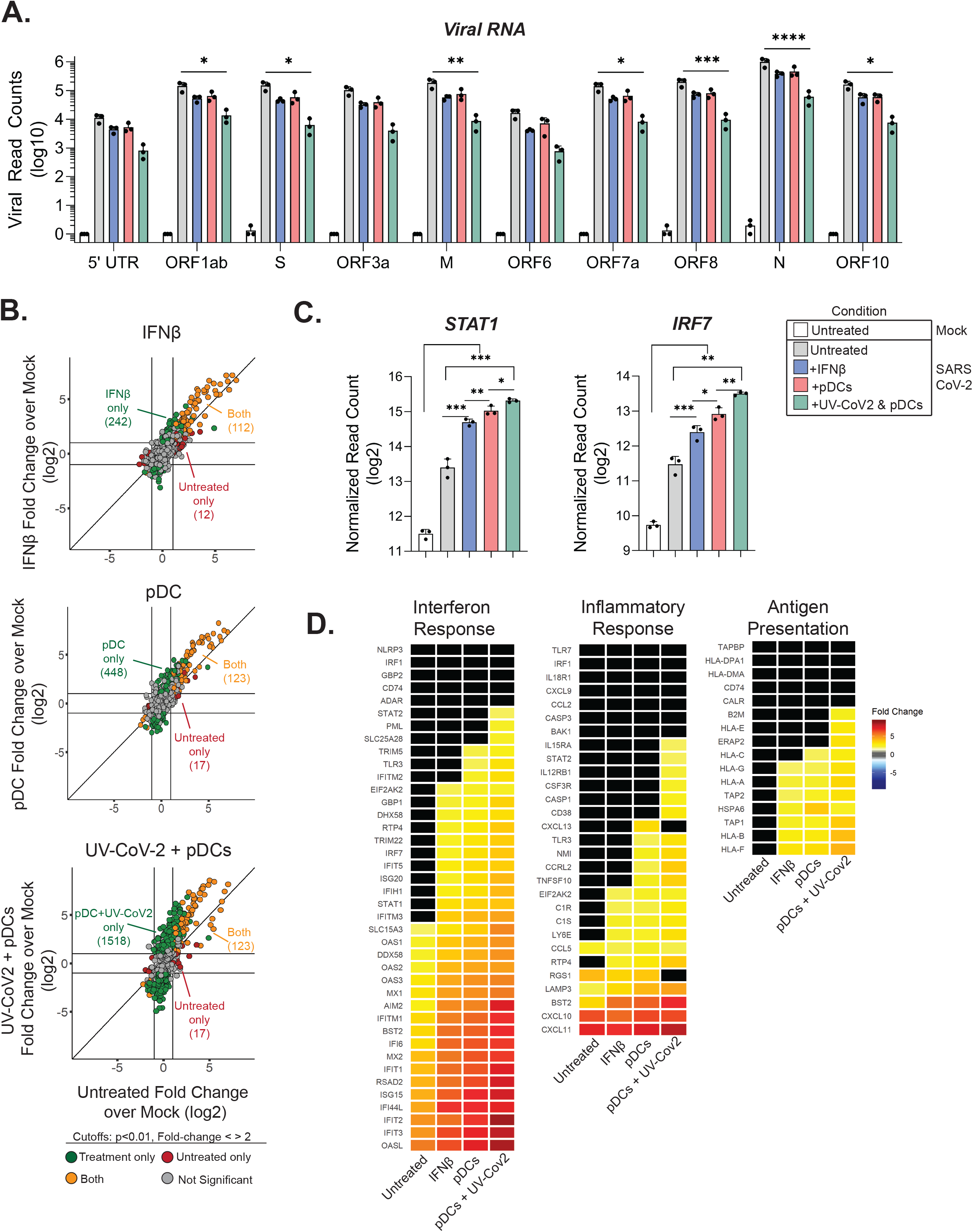
pDCs induce an interferon signature in pHAE cells. Bulk RNA-seq analysis was performed in pHAE cells from the co-culture experiment described in figure 3. A) Normalized read counts (log2) of SARS-CoV-2 RNA products, using the MT246667.1 reference sequence. B) Fold-change vs fold-change plot of SARS-CoV-2 infected pHAE cells vs uninfected pHAE cells and SARS-CoV-2 infected pHAE cells treated with IFN-β vs uninfected pHAE cells. Fold-change vs fold-change plot of SARS-CoV-2 infected pHAE cells vs uninfected pHAE cells and SARS-CoV-2 infected pHAE cells co-cultured with UV-SARS-CoV-2 treated pDCs vs uninfected pHAE and fold-change vs fold-change plot of SARS-CoV-2 infected pHAE cells vs uninfected pHAE cells and SARS-CoV-2 infected pHAE cells co-cultured with pDCs vs uninfected pHAE cells. Dots highlighted in green, red and orange are the DEG in each condition. C) Normalized read counts (log2) of expression of STAT1 and IRF7 in pHAE cells treated in the indicated conditions. D) Heatmap illustrating the z-scores for genes associated to the interferon response, inflammatory response and antigen presentation. Dots represent single replicates and bars represent the means. Data was analyzed using one-way ANOVA. *, P<0.05; **, P<0.01; ***, P<0.001.

To determine if the presence of activated pDCs in the co-culture changed the response of pHAE cells to infection, we plotted fold-change over mock for each co-culture condition (IFN-β treatment, pDC alone, UV-Cov-2+pDC) to the untreated infected control. Investigation of the differentially expressed genes (DEGs; cutoffs: p<0.01 fold change < >2) in infected pHAE cells treated with IFN-β treated cells or untreated identified that many DEGs were expressed in both conditions (112, yellow), but IFN-β treatment induced 242 uniquely expressed genes (green) while untreated cells only had 12 unique DEGs (red) (**Figure 4B**). Comparison of untreated pHAEs with pDC co-cultured pHAEs resulted in similar numbers of DEGs expressed by both or only untreated cells, however pDC co-cultured cells had 448 (green) unique DEGs. Co-culture with UV-CoV-2 stimulated pDCs resulted in the exact same number of DEGs expressed in both or only untreated cells, however there were three times more (1518 genes) genes uniquely expressed in co-cultured pHAEs when the pDCs were stimulated with UV-CoV-2 as compared to unstimulated pDCs (**Figure 4B**). This analysis determined that 1,518 genes were differentially expressed only if pHAEs cells were co-cultured with activated pDCs (**Figure 4B**). These results suggest that the presence of activated pDCs profoundly change the response of pHAEs to infection.

We next investigated the identity of the unique DEGs induced by co-culture of infected pHAE cells with pDCs. As expected, IFN-β induced expression of key signaling molecules and transcription factors in the type I IFN pathway such as STAT1 and IRF7. However, co-culture of pHAE cells with unstimulated pDCs or UV-CoV-2-stimulated pDCs had an even greater upregulation of STAT1 and IRF7 (Figure 4C). Accordingly, heatmap analysis identified an upregulation of many ISGs (IFIT2, RSAD2, ISG20) in IFN-β treated, pDC or UV-CoV-2-pDC co-cultured pHAE cells, with not only higher magnitude but also greater breadth of ISG expression in the pDC co-cultured cells (Figure 4D). We also analyzed if pDC co-culture affected the inflammatory response induced in pHAE by SARS-CoV2 infection. As seen in Figure 4D, co-culture with pDCs increased the inflammatory response of infected pHAE cells. Additionally, co-culture with UV-CoV-2-pDC upregulated the expression of 10 key inflammatory response genes, such as cytokine receptors IL12RB1, IL15RA, CSF3R, that was not observed in untreated, or IFNβ treated pHAE cells. Interestingly, co-culture with pDCs also induced the expression of molecules of the antigen presentation pathway. This may be of particular importance during the initial phases of infection in order harness the adaptive immune response to SARS-CoV-2.

## Discussion

In this study, we show that pDCs are able to recognize SARS-CoV-2 infection and produce high levels of type I IFN. Activated pDCs are able to control SARS-CoV-2 infection in pHAE cells by inducing an interferon response and an antiviral transcriptional program. SARS-CoV-2 and other coronaviruses are able to avoid recognition and type I IFN induction in several cell populations including airway epithelial cells. However, pDCs are able to sense SARS-CoV-2 and produce large amounts of type I and type III interferons. One major difference between pDCs and other cells is their expression of TLRs such as TLR7 which allows them to sense ssRNA^17^. Indeed, when we stimulated pDCs with UV-inactivated SARS-CoV-2 similar to live SARS-CoV-2, they produced IFN-α, suggesting that pDCs do not require active infection to sense the presence of SARS-CoV-2. A recent report has also shown that pDCs can get activated and upregulate CD80, CD86, CCR7 and OX40 after stimulation with SARS-CoV-2 independently of viral replication^36^. This gives pDCs two unique advantages for viral sensing: they are able to rapidly recognize the presence of viral RNA even before viral replication would start, without being susceptible to infection. Moreover, if pDCs don’t support productive SARS-CoV-2 replication, they may not be susceptible to the IFN antagonist effects of several SARS-CoV-2 nonstructural proteins.

We used human pHAE cell cultures to model the epithelial cell layer of the respiratory tract. pHAEs are susceptible to SARS-CoV-2 infection and recapitulate several features of respiratory disease, including inflammatory cytokine release and enrichment of ER stress pathways^24^. Co-culture of SARS-CoV-2 infected pHAEs together with pDCs that have been stimulated with UV-SARS-CoV-2 resulted in inhibition of viral particle production as seen by the reduction of viral titers in focus-forming assays (FFA) of the apical side of the pHAEs and inhibition of viral replication as seen in the reduction of viral RNA in the cells. Interestingly, co-culture of unstimulated pDCs with infected pHAE cells also resulted in reduced viral titers. We have previously demonstrated that viral particles are only released to the apical side of the pHAE culture^24^. Moreover, we confirmed the absence of infectious virus in the basolateral medium, where pDCs were located, by FFA. Nevertheless, pDCs were able to sense the infection of pHAE cells. Future studies will be needed to investigate if pDCs recognize viral RNA or soluble mediators from infected pHAE cells in the basolateral side.

The transcriptional analysis of SARS-CoV-2 infected pHAE cells corroborated that they are unable to produce type I IFN, although they can produce several inflammatory cytokines^24^. However, when infected pHAEs were co-cultured with activated pDCs, a set of 1510 additional genes were upregulated when compared to infected pHAEs that were not co-cultured with pDCs, revealing the strong impact of the presence of pDCs in pHAE cell response to viral infection. Several of these uniquely upregulated genes are ISGs or have been shown to have an antiviral role in the control of coronavirus infections, for example OAS or IFIT. Interestingly, the expression in pHAEs of genes upregulated by inflammatory cytokines increased by the presence of pDCs, suggesting that pDC production of inflammatory cytokines enhances the inflammatory response of pHAEs. We also observed the upregulation of genes involved in the antigen presentation pathway, which were not upregulated in the infected pHAE cells without pDCs. A key role of type I IFN signaling is the upregulation of class I antigen presentation to allow cytotoxic T cells to recognize viral infected cells and eliminate them. This is an additional mechanism by which pDC-derived IFN may contribute to viral control and induction of the adaptive immune response.

Since pDCs are able to sense SARS-CoV-2 infection and reduce viral replication, the remaining question is, how can SARS-CoV-2 overcome pDC recognition to establish infection. Previous studies have demonstrated that the frequency of pDCs in the blood of hospitalized Covid-19 patients is much lower than the frequencies in healthy donors^9^. This may be the result of three different mechanisms: pDCs may be activated very early on during infection, and in individuals where infection is more severe we observe the loss of the previously activated pDCs. A study by Swiecki et al. showed that pDCs die by apoptosis after viral recognition and IFN-α production^37^. Consistently, we observed an increase in the apoptosis signature in SARS-CoV-2 stimulated pDCs. Moreover, it has been also shown that during prolonged or chronic viral infections, pDCs present an exhausted phenotype and stop producing type I IFN^38,39^. Another possibility is that SARS-CoV-2 viral proteins directly target pDCs *in vivo* and result in their depletion. Network analyses of single-cells RNA seq of COVID-19 patients showed an association between apoptosis in pDCs and disease severity^22^. Finally, it is also possible that pDCs are largely recruited to the infected tissues such as lungs and this results in reduced frequencies in peripheral blood. Indeed, numbers of pDCs are found in the BAL of mild COVID-19 patients in contrast to severe COVID-19 patients which have reduced numbers of pDCs in BAL^10^. Future studies in animal models, where the initial hours after infection can be studied will be very helpful in elucidating the fate of pDCs and why they are reduced in COVID-19 patients. Recently, a study in which Covid-19 patients were treated with IFN showed that early interferon treatment associates with favorable clinical response in COVID-19 patients^15^. Here we showed that co-culture of pDCs with infected pHAEs was better at controlling SARS-CoV-2 infection than pre-treatment of pHAEs with IFN-β, suggesting a potential use of pDC activation as treatment for COVID-19. Since pDCs are able to produce not only IFN-β, but IFN-α, and IFN-λ, rescuing pDC numbers or activating the remaining pDCs in COVID-19 patients is an attractive strategy to reduce the severity of the disease.

## Author contributions

L. C-B., and M.S. contributed to the acquisition, analysis, and interpretation of the data, as well as the conception and design of the study, and wrote the manuscript A.V. contributed to the acquisition, analysis, and interpretation of the data, and wrote the manuscript. C.J.R., P.R, M.D-G., and T.C. contributed to the acquisition and analysis of the data, L.J.A. and A.G. contributed to the interpretation of the data.

